# Optical and physical mapping with local finishing enables megabase-scale resolution of agronomically important regions in the wheat genome

**DOI:** 10.1101/363465

**Authors:** G. Keeble-Gagnère, P. Rigault, J. Tibbits, R. Pasam, M. Hayden, K. Forrest, Z. Frenkel, A. Korol, E. Huang, C. Cavanagh, J. Taylor, M. Abrouk, A. Sharpe, D. Konkin, P. Sourdille, B. Darrier, F. Choulet, A. Bernard, S. Rochfort, AM. Dimech, N. Watson-Haigh, U. Baumann, P. Eckermann, D. Fleury, A. Juhasz, S. Boisvert, M-A. Nolin, J. Doležel, H. Šimková, H. Toegelová, Jan Šafář, M-C. Luo, F. Camara, M. Pfeifer, D. Isdale, J. Nystrom-Persson, IWGSC, D-H Koo, M. Tinning, D. Cui, Z. Ru, R. Appels

## Abstract

**Background:** Numerous scaffold-level sequences for wheat are now being released and, in this context, we report on a strategy for improving the overall assembly to a level comparable to that of the human genome.

**Results:** Using chromosome 7A of wheat as a model, sequence-finished megabase scale sections of this chromosome were established by combining a new independent assembly based on a BAC-based physical map, BAC pool paired end sequencing, chromosome arm specific mate-pair sequencing and Bionano optical mapping with the IWGSC RefSeq v1.0 sequence and its underlying raw data. The combined assembly results in 18 super-scaffolds across the chromosome. The value of finished genome regions is demonstrated for two approximately 2.5 Mb regions associated with yield and the grain quality phenotype of fructan carbohydrate grain levels. In addition, the 50 Mb centromere region analysis incorporates cytological data highlighting the importance of non-sequence data in the assembly of this complex genome region.

**Conclusions:** Sufficient genome sequence information is shown to be now available for the wheat community to produce sequence-finished releases of each chromosome of the reference genome. The high-level completion identified that an array of seven fructosyl transferase genes underpins grain quality and yield attributes are affected by five f-box-only-protein-ubiquitin ligase domain and four root-specific lipid transfer domain genes. The completed sequence also includes the centromere.

## Background

The hexaploid wheat genome has been assembled into 21 pseudomolecules that cover more than 90% of the estimated 15.7 Gb of DNA that constitutes the genome (1). Unlike previous efforts to sequence the wheat genome (2, 3, 4), the IWGSC RefSeq v1.0 assembly of pseudomolecules provides a high-quality linear assembly of each chromosome from one terminal region through the centromere to the other terminal region in the form of 70-80 super-scaffolds per chromosome. Unlike advanced assemblies of human, and model organisms (5), which all included sequencing of BAC-based physical assemblies, the IWGSC RefSeq v1.0 assembly was achieved by combining a primarily whole-genome short read based assembly with Hi-C, BAC sequencing and genetic/optical mapping information. The algorithmic advances that have made the IWGSC RefSeq v1.0 assembly possible leave a final challenge of bringing the local base level assembly up to a finished status, where the assembly is contiguous at the megabase scale, with no gaps (Ns).

The drive for finishing the human genome has come from the requirement that all genes should be accounted for in order to establish complete coverage for functional studies (6, 7). In the same way, a finished genome for wheat is required to understand the dynamic nature of the wheat genome (2, 8), its capacity to adapt to hot and dry environments as well as very cold and wet regions, and to capture genes responsible for traits such as yield, salinity tolerance, faster germination time, or nutritional quality for fundamental and translational research. The capacity to adapt and produce grain for a variety of food and non-food products accounts for the prominent position of wheat in the modern industrial supply chain (9). The gene space for chromosome 7A was partially defined by the IWGSC Chromosome Survey Sequence (CSS) assembly (2) and codes for genes involved in determining the quality of flour (seed storage proteins, enzymes for starch and fructan synthesis, yellow pigment, preharvest sprouting tolerance) as well as many abiotic responses. Yield is widely acknowledged to be a complex trait and components that are considered to be stable contributors to this trait include thousand kernel weight (11) and spikelets per spike (12, 13), both having significant associations with a region on 7A (13, 14, 15). Other trait components contributing to grain yield such grains per spike and vernalisation requirements, as discussed in (5,10), are also located to the same region on chromosome 7A and together these define an important candidate target region for finishing. Another region coding for a major contribution to grain quality (grain fructan content, 16), provides a second target region. In the assembly reported in the present paper, the centromere, generally considered one of the most challenging regions of the genome to assemble, was also considered using Bionano (17) maps to both confirm the assembly and to provide direction for resolving inconsistency between cytological and assembly data. Manual annotation was performed based on the automated annotations (1)(RefSeq annotation v1.1), using alignments of available RNA-Seq data (3, 18) to ensure gene models were consistent with transcriptome evidence.

In the present study, we used the GYDLE bioinformatics (https://www.GYDLE.com/) software suite to produce an independent assembly of chromosome 7A which integrated a new BAC-based assembly, high resolution genetic and Bionano map assemblies, as well as chromosome specific mate-pair data and BAC-based physical maps. We then demonstrate the feasibility of finishing targeted regions including agronomically important regions of chromosome 7A by using the GYDLE tool suite (https://www.GYDLE.com/) to simultaneously assess and combine our assembly with the IWGSC RefSeq v1.0 assembly in an iterative process that re-used available raw data to resolve inconsistencies between assemblies, and between assemblies and the raw data. This approach highlights that simultaneous use of sequence and mapping resources generated by different technology platforms allows greater progress towards complete resolution of genome sequences than otherwise possible by using individual technologies. It is the first true demonstration of independent genome assembly integration that is not based on a facile merge overlap process and provides a tractable route for finishing almost any genome region of interest in wheat, or in fact the whole wheat genome if applied universally.

## Results

### BAC and optical map-based assembly of chromosome 7A

We assembled chromosome 7A of hexaploid wheat into 72 islands (defined below) covering a total of 752 Mb of DNA. The assembly combined a range of data sources including a 755 Mb physical map comprising 732 BAC contigs, represented by 11,451 BACs in 732 minimum tiling path (MTP) BAC sets, as well as mate-pair sequencing of genome-wide and chromosome-arm specific libraries (Methods and Additional file 1) and chromosome-arm specific Bionano optical maps.

The islands are the combined result of scaffolding the individual BAC pool assemblies (which total 711 Mb of sequence in 4,107 sequence contigs) using both Bionano maps (546 maps covering 746 Mb) and sequence alignments. The largest island covers 59.9 Mb and 71% of the assembly is represented by 20 islands larger than 10 Mb. Our sequence assembly is highly contiguous locally with a contig mean length of 173Kb and 95% (678.6 Mb) of its total length in 789 contigs over 100Kb. Very high base-level accuracy and sequence continuity was achieved through the simultaneous integration of both BAC pool and mate-pair sequencing data, physical mapping information and Bionano alignments (Fig. 1).

**Fig. 1.**
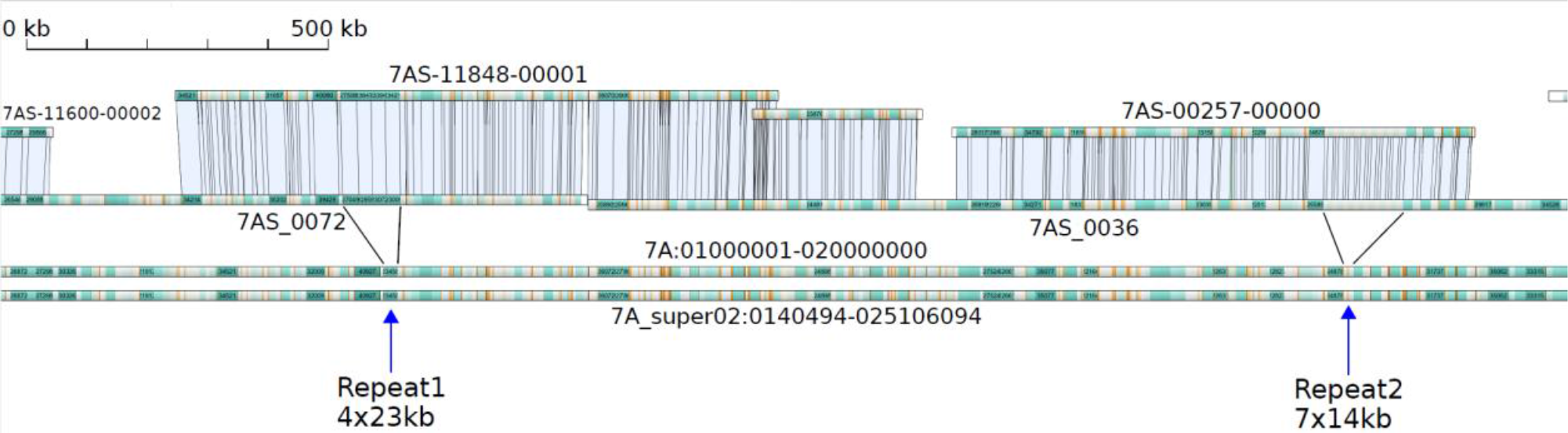
GYDLE assembly (top tracks) aligned to the IWGSC RefSeq v1.0 chromosome 7A pseudomolecule (bottom tracks, see (1)). The top two tracks show BAC pools 7AS-11848, 7AS-11877 and 7AS-00257 aligned to Bionano maps 7AS_0072 and 7AS_0036. The BAC pool assemblies are finished with no gaps or ambiguities and have resolved repeat arrays which are collapsed in the IWGSC RefSeq v1.0 assembly. Depending on the coverage of BACs, regions of the IWGSC RefSeq v1.0 assembly are either covered by a single BAC pool, covered by multiple BAC pools (such as the 30Kb of overlap between 7AS-11848 and 7AS-11877) or not covered by any BAC pool (such as between 7AS-11877 and 7AS-00257). The GYDLE assembly increased the assembled sequence length by a total of 169Kb across the region covered by these three pools (approximately 8%).

### Physical map assembly

Flow-sorted 7AS and 7AL telosomes (Additional file 1) were used to construct BAC libraries, comprising 58,368 and 61,056 clones respectively, which were all fingerprinted as described in (19). LTC software (20) was used to establish ordered assemblies of the BAC clones from the DNA fingerprint information in order to select a minimum tiling path (MTP) BAC set for sequencing (Additional file Figure S3). The following procedure was carried out for 7AS and 7AL independently: a network of “overlaps” was constructed using Sulston score cutoff 1E-10. Five iterations of increased stringency were applied in increments of 1e-5, as well as Q-clones being identified at each step (20). BAC contigs with less than 5 clones were not included in the final outputs. The physical assembly comprised 380 BAC contigs on 7AS (352 on 7AL) and contained 42,244 clones on 7AS (47,776 on 7AL), from which an MTP of 5,280 clones for 7AS (5,832 on 7AL) was defined with estimated total length for 7AS of 353Mb (402Mb for 7AL).

### BAC set assemblies

The 732 MTP BAC sets were sequenced in 813 pools, with each pool comprising no more than 40 BACs (the median number of BACs per pool was 11). This generated 1.67 billion paired reads, which were first assembled independently for each BAC set using ABySS (21) to produce a stage 1 assembly of 882 Mb of sequence in 74,572 contigs. These contigs were used to seed a stage 2 assembly based on the use of Nuclear, Resolve and Vision software (https://www.GYDLE.com/). These tools allow the sensitive alignment of raw data, resolution of conflicts with raw data, together with real-time visualization, to assemble BAC sets simultaneously using all available datasets. The datasets included the BAC set paired-end reads, mate-pair reads from whole genome and flow-sorted 7AS and 7AL telosomes, and the raw data from the 7AS and 7AL survey sequencing (2). This hybrid assembly further used physical mapping information (BAC-end derived reads identified using the cloning vector, raw fingerprinting data and BAC ordering) to produce assemblies consistent with the MTP layout along BAC sets and to identify and quarantine contaminant BACs for separate assembly and placement. As part of stage 3, multiple rounds of automated contig correction, extension and scaffolding, with manual curation in target regions, produced 1,897 scaffolds for 7AS (2,211 for 7AL).

### Bionano map assembly and island construction

Bionano optical data were generated from independently flow-sorted 7AS and 7AL telosomes producing 360,390 molecules on 7AS (416,563 on 7AL), representing 192x coverage on 7AS (238x on 7AL). The Bionano IrysView software was used to assemble the 178,217 7AS molecules into 783 optical maps (145,207 molecules into 330 maps for 7AL). The total length of the optical maps was 447 Mb for 7AS (413 Mb for 7AL) with an N50 length of 1.55Mb on 7AS (2.07 Mb on 7AL). These data and the BAC set stage 2 scaffolds were combined using GYDLE optical mapping and assembly software to produce islands, representing connected sets of sequence scaffolds and optical maps. This process included a map validation step using molecule alignments to identify a set of high confidence maps (272 maps on 7AS, 270 on 7AL), and the improvement of BAC set assemblies by using optical alignments for stitching, orienting and locally polishing scaffolds. This produced 72 final islands covering 752 Mb, 711 Mb of which was covered by BAC set sequences in 4,107 contigs.

### BAC set finishing and assembly integration

Several regions of the chromosome were selected for designing our finishing process (stage 3), using the GYDLE software with an emphasis on complete data integration and systematic human visual review in order to achieve BAC set assembly completion: namely, a single, gapless contig of finished-quality sequence per BAC set supported by the consistency of sequence, physical mapping and optical data at the raw and assembled level, including the resolution of close repeats. We finished 30 BAC sets (representing 25 Mb) with this process and extended it to allow the inclusion of IWGSC (assembly and raw) data to compare, qualify and integrate the assemblies, with the view to being able to finish a sequence for the whole chromosome (i.e outside BAC sets as well).

### Overview of GYDLE and IWGSC RefSeq v1.0 chromosome 7A assemblies

The IWGSC RefSeq v1.0 assembly of chromosome 7A represents 736.7 Mb (~90.4%) of sequence distributed relatively uniformly across the chromosome. A major strength of the IWGSC RefSeq v1.0 is the long-range organisation of scaffolds and super-scaffolds into pseudomolecules. The chromosome 7A scaffolds are made up of 27,657 contigs, with a mean length of 26.2 Kb, and 11.7 Mb of unresolved bases (N) in sized gaps, internal to scaffolds. Hence the IWGSC RefSeq v1.0 has a representation of most of the chromosome 7A order and arrangement but with many small gaps internal to scaffolds, and a smaller number of large gaps of unknown size between scaffolds (linkage evidence but no gap size estimation) and between super-scaffolds (no linking evidence). Our GYDLE assembly represents 752 Mb of the 7A chromosome, with 711 Mb in near-complete assemblies of the BAC sets, which are ordered and oriented into islands with larger (mostly sized) gaps between BAC sets. Fig. 1 highlights the structural differences between the assemblies, showing the near-complete representation of the underlying sequence and the concordance with Bionano optical maps within BAC pools in our assembly and the gaps between them often filled with IWGSC RefSeq v1.0 sequence. Fig. 1 also highlights that in this case the GYDLE assembly correctly represents the number of large tandem repeat sequences which are collapsed in the IWGSC RefSeq v1.0. These repeats are documented by Bionano maps and add about 8% to the total length of the region. This observation is consistent with the IWGSC RefSeq v1 wheat genome (1) which argued that much of the missing genome length in the assembly was from under-representation of arrays of repetitive sequence units.

### Classifying chromosome 7A into 18 connected components

Super-scaffolds define the extent of sequences which are internally connected, ordered and in most cases oriented through underlying data links (physical or Bionano maps) without necessarily establishing the complete sequence in between or gap size. The 35 super-scaffolds of the IWGSC RefSeq v1.0 chromosome 7A pseudomolecule (1) were constructed using Hi-C ordered scaffolds, with scaffold joins made where either the physical map via Bayer-IWGSC Whole Genome Profiling (WGP^TM^) tags (1) or Bionano maps provided a link – a process that is sometimes prone to error due to the repetitive nature of sequences that occur at the end of scaffolds in the IWGSC RefSeq v1.0 assembly. Our island assembly integrated the physical map and Bionano data with underlying sequence enabling further and more accurate super-scaffolding.

Using our island-assembly we were able to reduce the 35 super-scaffolds in IWGSC RefSeq v1.0 to 18. Our assembly could also orient the remaining two IWGSC RefSeq v1.0 scaffolds (of 193) that were un-oriented in chromosome 7A (7AS-00257-00000 orients scaffold138751 in minus orientation; 7AS-12029-00000 orients scaffold17971 in minus orientation). This completes the scaffold orientation across the whole of chromosome 7A. Our 18 super-scaffolds were aligned to a new high density genetic map calculated from assigning over 4,000 markers to 900 progeny genotyped by Genotype-By-Sequencing (GBS), from an 8-way MAGIC cross integrated with the bi-parental Chinese Spring x Renan genetic map for chromosome 7A (Fig. 2A; Additional file 2a, 2b), and this supported the overall super-scaffold order and orientation.

**Fig. 2.**
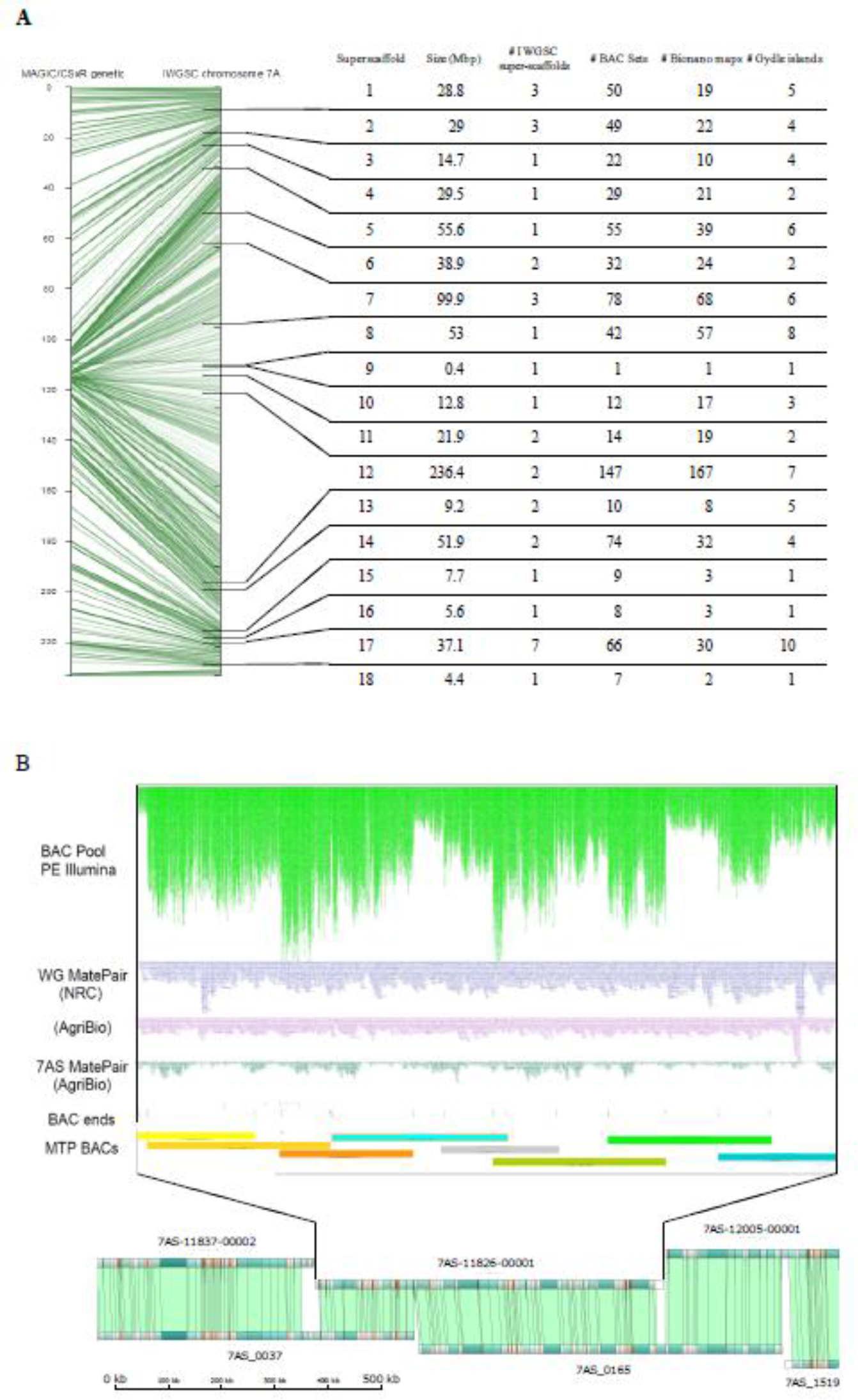
A) Alignment of MAGIC/CSxRenan genetic map (left axis, Additional file 2b) against IWGSC RefSeq v1.0 chromosome 7A (right axis). On the right axis, ticks denote the boundaries of the 18 super-scaffolds defined in this manuscript. The table summarises the assembly information integrated in each super-scaffold (see also Additional files 4b, 5). Some cross-overs in the alignment of the MAGIC and IWGSC genetic maps reflect ambiguities that can arise as a result of the high and distributed repetitive sequence content of the wheat genome combined with the fact that the MAGIC map is based on a multiple cross between 8 modern varieties and the physical map is Chinese Spring. In some cases the map suggested no linkage between markers located in a physical contig. If reexamination of the physical contig indicated a ‘weak link’ in the physical contig assembly (example shown in Additional file, Fig. S3) then the assembly was split into ‘a’ and ‘b’ contigs. If the physical contig evidence was unambiguous, the markers were set aside for reconsideration in light of more evidence being obtained. B) An example of a locally finished sequence (BAC pool 7AS-11826; 655Kb) showing integration of multiple data types: paired-end Illumina data from BACs (top, green); three independent mate-pair libraries; Minimum tiling path (MTP) BAC start and end points, based on mapping junction with vector; Bionano optical map alignments. Note that coverage of BAC pool data varies depending on double- and triple-coverage of BACs in MTP. Sequence is contiguous with no gaps. The assembled sequence joined two Bionano maps. This 655 Kb contig, which included the P450 gene, TaCYP78A3, shown to be associated with variation in grain size (33).

Using Nuclear (software, see Methods and Additional file 3) to align IWGSC RefSeq v1.0 contigs (27,651 contigs; length: 724.64 Mb) to the GYDLE assembly v3.0. and stringent mapping parameters we found 11,050 contigs matched the GYDLE assembly exactly (193.35 Mb), 13,539 contigs that had a partial (>90%) sequence match (484.54 Mb), while the remaining 3,062 contigs (46.75 Mb) had no matching sequence in the GYDLE assembly (consistent with the missing sequence between BAC sets). Using a stringent approach these alignments were used to identify potential gap sequences, where only gaps between consecutive mappings of IWGSC RefSeq v1.0 contigs within GYDLE contigs were selected both within scaffold and between scaffold gaps. We were able to bridge 82 of the 193 scaffold-scaffold gaps in the IWGSC RefSeq v1.0 assembly with GYDLE assembly contigs spanning IWGSC v1.0 inter-scaffold gaps. Of these, 26 had a clean mapping of the flanking IWGSC RefSeq v1.0 contigs suggesting consistency between assemblies for these regions. The reduction of 82 to 26 bridging locations reflects the a priori difficulty expected with these scaffold-scaffold sequences and our conservative approach with the edges of scaffold assemblies in the IWGSC RefSeq v1.0 often conflicting with the GYDLE assemblies. For comparison, the same analysis with the Triticum 3.0 (subsequently referred to as PacBio) assembly (4) found 88 scaffold-scaffold gaps bridged, with 54 of these in common with the GYDLE set, though in only one case were the GYDLE and PacBio bridging sequences the same length (Additional files 4a, 4b). These scaffold-scaffold gaps are clearly tractable although they will require careful resolution, preferably combining other assembly information before bridge sequences can be determined across the wheat genome. For intra-scaffold contig-contig gaps we identified 3,016 contig mappings with perfect flanking contig alignments to the GYDLE assembly (Additional file 5). In total the contig-contig gap filling replaced gap of Ns with 562,594 bp of sequence, with a mean gap size of 152.6 bp among the 2,655 non-zero length gaps. The contig-contig gap sequences were observed to be generally either GC rich, often containing long homopolymer G or C runs, or contained di-and tri-nucleotide (and higher order) repeat sequences. Unanchored IWGSC RefSeq v1.0 scaffolds could also be assigned to chromosome 7A and accounted for 19.4 Mb of un-scaffolded sequence being identifiable as 7A against our assembly.

To assess the gene-level agreement between assemblies, we extracted the respective genome sequences (from the beginning of 5’ UTR to end of 3’ UTR) from the IWGSC RefSeq v1.0 annotation for chromosome 7A and used these to query the GYDLE sequence. We found 13,283 (96.1%) genes were present in the GYDLE assembly; of these 11,312 (81.8%; 4,370 high confidence (HC) and 6,942 low confidence (LC) (76.6% and 85.4% of their respective totals) genes matched perfectly to IWGSC sequences. Of the non-perfect matches, 414 (3%) matched across the full length but with base-pair mismatches; 1,557 (11.3%) did not match across their full length. Across chromosome 7A we identified 107 (54 HC and 53 LC) genes in the IWGSC RefSeq v1.0 annotation which contained gaps (stretches of Ns) in the coding sequence (Additional file 6). Of these, 100 were complete in our GYDLE sequence.

### Local finishing of a genome region associated with grain fructan content

We identified a tight cluster of markers on chromosome 7A associated with grain fructan levels in a GWAS analysis of 900 wheat lines using NMR (3.8 ppm proton shift, see Additional file 7) and genome-wide SNP markers (derived from exome capture assays). The markers were contained in a single BAC contig 7AS-11582 within a 7.5Mb island (Fig. 3), corresponding to the IWGSC RefSeq v1.0 region spanning 3,070,713bp to 5,459,064bp. The 7AS-11582 contig was targeted for finishing. The tandem repeated element (four units of a 10 Kb repeat sequence; Bionano map, Fig. 3B), was sequenced using a single BAC (7AS-066B03) covering that repeat and PacBio sequencing combined with short-read Illumina data, physical mapping and optical data during the finishing process.

**Fig. 3.**
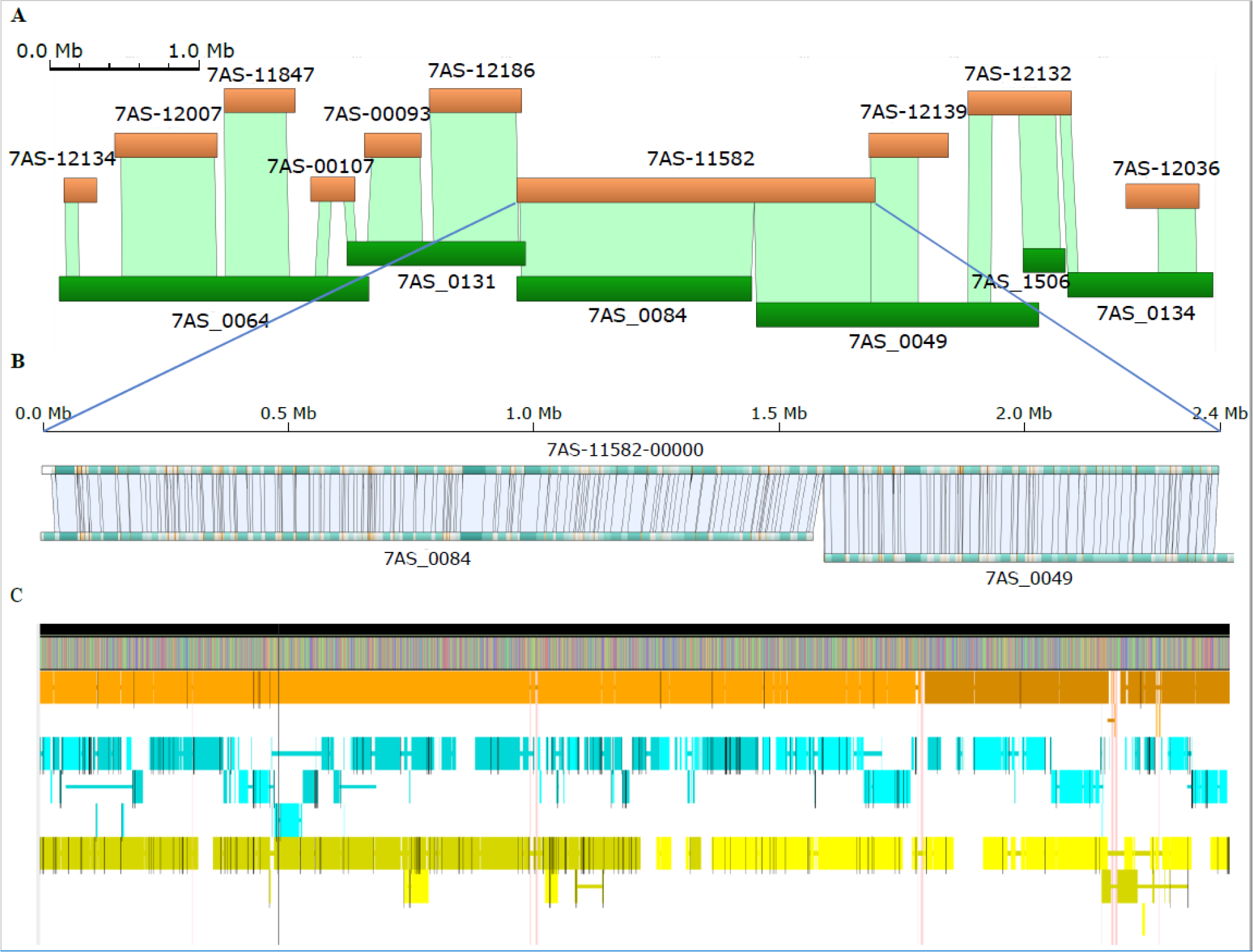
Detail of local region associated with fructan content. A) The 7AS island containing 7AS-11582. B) Optical maps (7AS-0064 and 7AS-0049) aligned against the finished sequence for 7AS-11582. C) Finished GYDLE sequence for 7AS-11582 (top) with alignments of matching contigs/scaffolds from IWGSC RefSeq v1.0 (orange), TGAC (cyan) and PacBio (yellow) assemblies (see Supplementary Methods). Gaps are indicated by white space between HSPs and differences by black bars. Vertical pink links indicate regions of the finished sequence not present in any other assembly.

Through iterative rounds of automated and manual assembly we constructed a final sequence assembly, integrating and consistent with all available raw data sources, of 2,397,551 bp in a single contig with no gaps or unresolved bases. The IWGSC RefSeq v1.0 sequence comprised 3 scaffolds and 105 internal gaps (giving a total of 107 gaps across the region, Additional file 8, Figure S5). Of these, 98 were filled with sequence from the GYDLE assembly, with a common observation that the gap sequences were either GC rich (12 gaps with 80%+ GC) and/or contained a homopolymer run of G10+ or C10+ (52 gaps). Illumina sequencing technologies are known to have difficulties in regions with G/C homopolymer runs (22) and, while the coverage in these regions is generally lower than that of surrounding sequences, supporting raw data for these missing sequences is often present in existing data sets. The longest filled gap sequence in the 7AS-11582 region was 6,826 bp with a mean filled gap size of 306 bp (median 92 bp). The remaining seven gaps were closed with either short sequence overlaps between neighboring contigs or subtle rearrangements of the final sequence versus the original contig order. A small number of within-contig insertions (eight) and deletions (nine) were also found. The majority of these were single bases and three were greater than 100 bp.

We identified scaffolds from the TGAC (3), PacBio (4) and IWGSC RefSeq v1.0 (1) assemblies using our finished sequence as bait and applying the same alignment parameters for each. Fig. 3C summarizes these assemblies aligned against the finished GYDLE 7AS-11582 sequence. As expected, no assembly fully represented the entire region and all assemblies were fragmented around the repeats, highlighting their difficulty for assembly. Comparison of assembly completeness and similarity across this region indicates that assembly merging as a means of genome finishing will require a careful strategy capable of deciding between competing options. Clearly, simple merge-overlap approaches are not likely to improve the entire genome representation provided in the IWGSC RefSeq v1.0 and that an approach that re-references the raw data (preferably from multiple sources simultaneously) to resolve inconsistencies will be required.

One of the most important attributes of having a locally finished sequence is the impact on the accuracy of the gene annotation. There were 62 HC and 68 LC genes annotated across the 7AS-11582 region. Five of the HC had gaps within the genomic sequence and of these, two (TraesCS7A01G010500 and TraesCS7A01G010800) had gaps within their CDS. The finished assembly completed these genes and enabled the gene models to be updated. For TraesCS7A01G010500 the gene model was incomplete in all other available annotations of wheat and the finished gene model was found to be a novel variant of a BAG family molecular chaperonin regulator seven gene (UniProtKB - Q9LVA0 (BAG7_ARATH). Close proximal regions to genes generally harbor functional elements and the finishing process in these regions closed 38 (18 HC; 20 LC) gaps within 5 Kb of annotated genes. Of particular interest for grain quality was the identification, confirmed through the manual curation of the gene models across the finished sequence, of a tandem array of seven glycoside hydrolases (EC 3.2.1, labelled a to g), including the gene model GH32b being assigned as a 1-FFT (fructan 1-fructosyltransferase) on a sequence similarity basis and GH32g to 6-SFT (sucrose:fructan 6-fructosyltransferase). Both these genes are expressed in the grain and stem, based on alignments of RNA-Seq data from (18) and represent good candidate genes for variation in grain fructan levels.

### Local finishing of a genome region associated with grain number and weight

Published studies have mapped yield QTL to the long arm of chromosome 7A with varying degrees of resolution (23). Using a RAC875xKukri cross, we mapped yield and two yield components, thousand kernel weight (TKW) and kernels per spikelet across the length of chromosome 7A (Additional files 7 and 9). A cluster of four TKW QTL was in the 172.4-177.0 cM region of the RAC875/Kukri map (Additional file 9). These co-located with the QTL TaTKW-7AL which was mapped to a 1.33 cM interval on chromosome 7A (between 90K SNPs IWB13913 and IWA5913; (15)) and a QTL for spikelet number per spike (13) in the same interval. These QTL define a core yield QTL region located between 672,014,054 bp and 674,276,807 bp in the IWGSC RefSeq v1.0 7A pseudomolecule, which we targeted for complete sequence finishing (Fig. 4). The region is covered by two scaffolds (scaffold274331-1 and scaffold91613) in IWGSC RefSeq v1.0 (1), where the 2.262 Mb pseudomolecule sequence contains 37,065 uncalled bases (N) in 101 gaps. In the GYDLE assembly, the core region, contained within a single island, was covered at 94% by 5 non-overlapping BAC sets (7AL-12138, 7AL-05057, 7AL-12241, 7AL-00419 and 7AL-11456). We performed finishing on these BAC sets to produce 2,125,896 bp of the region, then finished the intervals between BAC sets using the raw sequence data (IWGSC and our 7A mate-pair libraries) combined with Bionano to resolve 144,424 bp. The finished core yield QTL region is a gapless contig of 2,270,131 bp (Additional file 10).

**Fig. 4.**
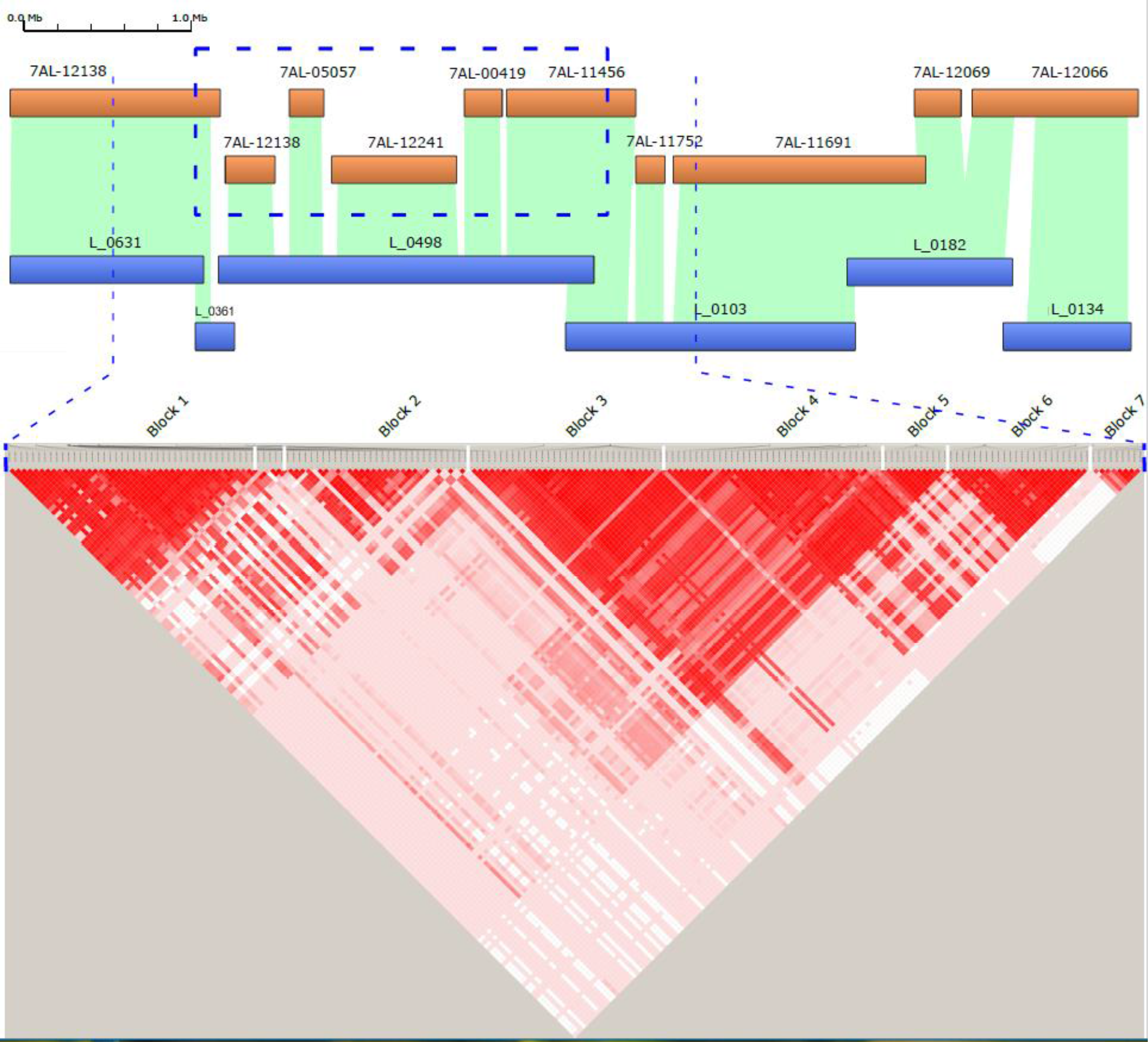
GYDLE island containing the core yield region (defined by blue dotted lines, coordinates 671200000-675300000 bp). Assembled GYDLE stage 2 sequences (orange, stage 2 with the genome segments based on BAC pools) aligned to Bionano maps (horizontal blue bars) in the top panel. The genome sequence within the bold dotted blue box in the top panel is the stage 3, finished, genome sequence region. The lower panel displays pairwise LD values (D’, (42)) between a total of 203 gene-based SNPs in same region across 863 diverse bread wheat accessions. Only common SNPs with high minor allele frequency (MAF > 0.3) are shown because common SNPs have high ability to define extent of LD and historical recombination patterns in diverse collections. The SNPs present within 2000 bp on either side of gene were included in this analysis. Color code: Bright red D’ =1.0 and LOD ≥ 2.0 (high LD); light shades of red indicate D’ < 1.0 and LOD ≥ 2.0 (low-medium LD); white indicates D’ <1.0 and LOD <2.0 (no LD or complete decay).

Manual curation of all the IWGSC gene models across this region enabled many small annotation inconsistencies to be detected and corrected, most of which arose because of micro-assembly ambiguities. Across the QTL core region there were 61 genes (27 HC and 34 LC) annotated in IWGSC V1.0 of which 6 had gaps within their genomic sequence in the original assembly (Additional file 11). The sequence downstream of the core QTL (674,273,097 to 674,876,866 bp) contained 27 annotated genes (12 HC and 15 LC) which included a cluster of 8 Hydrophobic-domain protein family genes ((1), cortical cell delineating class, specifically expressed in roots). We used the available finished sequences to investigate linkage disequilibrium across the QTL region in 863 unrelated wheat accessions each assayed with Roche exome capture technology (Fig. 4; Additional file 7). Seven blocks of high LD are seen across the region and clearly define targets for further fine mapping of the thousand-kernel weight (TKW) and kernels per spikelet in blocks 2-5 (Fig. 4). The gene function predictions based on the domains in the translated protein sequences (Additional file 11) serve to further refine a candidate gene list.

### Multiple windows into the wheat chromosome 7A centromere

Centromeres mediate chromosome attachment to microtubules and ensure proper segregation of the sister chromatids during mitosis and meiosis (24). While the active centromere and associated kinetochore complex is characterized in plants by the location of CENH3 binding sequences (25) various working definitions include reduced recombination rates, methylation patterns, transposable element repeat patterns and constitution, and chromosomal centromere-break points. Taking a classical definition of the centromere as the region of suppressed recombination, we defined a centromere region in chromosome 7A based on an analysis of over 900 lines in an 8-way MAGIC population cross, genotyped with a targeted GBS assay (Additional file 2), and determining the parental donor of chromosome segments for counting cross-overs. The centromere region defined by suppressed recombination spans almost half the chromosome, between approximately 175Mb-600Mb (425 Mb). Within this region a 170 Mb (spanning 270-440 Mb) region of no cross-overs containing a smaller 60 Mb region (spanning 320 Mb-380 Mb) enriched for centromere-specific CRW (Cereba/Quinta) repeat families was identified (Fig. 5A).

**Fig. 5.**
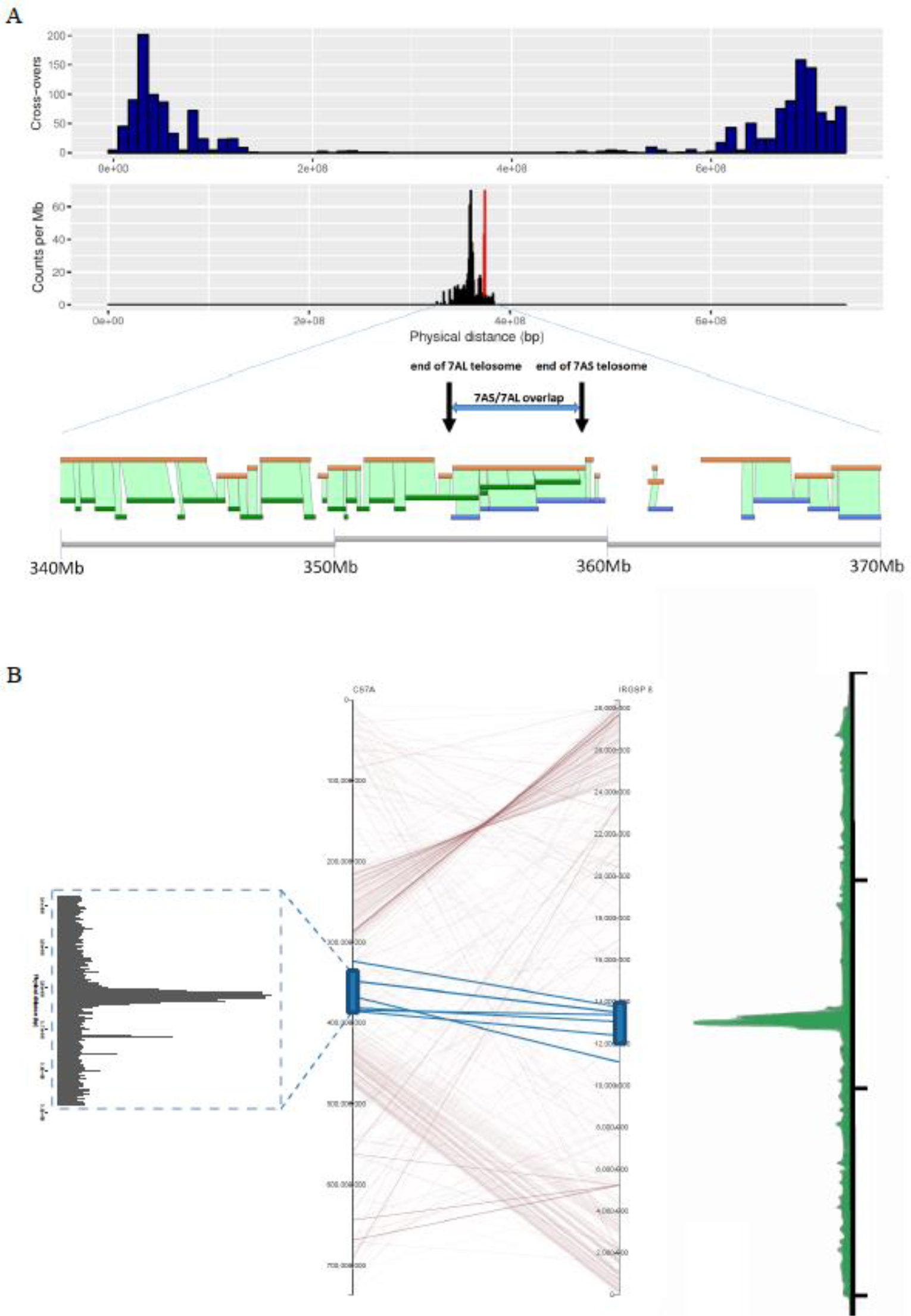
A) The 7A centromere. The top panel shows cross-over counts from an analysis of 900 lines (only cross-overs from 465 lines shown; see Additional file 1) of a MAGIC population (10 Mb bin size) across the entire chromosome and identifies a region of zero recombination traditionally associated with the centromere. The second panel shows this region is the primary location of the Cereba TEs that define wheat centromeres. Within this region we also identified a compact cluster of Tai 1 sequence elements shown in red. The third panel indicates the location of the breakpoints that generated the 7AS and 7AL telosomes and the bottom panel shows the GYDLE islands (sequences in orange) and Bionano maps (7AS in green, 7AL in blue) for this region tiling the IWGSC RefSeq v1.0 (grey) from 340Mb to 370Mb. The break in both the GYDLE and Bionano maps in the 349 Mb region is referenced in the text as well as Fig. 6A as a possible location of CENH3 binding sites. B) The 7A centromere aligned to rice chromosome 8. Lines indicate syntenic genes, with conserved gene models between the two centromere regions highlighted in blue. Equivalent locations of the CENH3 binding sequences shown on the right and left sides. The CENH3 plot for the rice 8 centromere (right side) was modified from Yan et al (34).

Alignment and anchoring of the broad centromere region defined by the CRW sequences to the rice chromosome 8 functional centromere region (Fig. 5B) identified six highly conserved genes (TraesCS7A01G284500, TraesCS7A01G288700, TraesCS7A01G294900, TraesCS7A01G295300, TraesCS7A01G295400, TraesCS7A01G292400) that spanned both the 7A and rice 8 centromere regions and were homologs to the rice genes identified by Yan et al (2008) as highly conserved across crop plants. Manual annotation of 7A genes following gap closure allowed the functional domain of the 7A centromere to be defined through synteny alignment (Fig. 5B, Additional file 12) to the rice chromosome 8 centromere.

Complete agreement between separate 7AS and 7AL telosome assemblies and data (raw flow-sorted chromosome paired-end read data (2), GYDLE BAC sets and Bionano maps sequences) provided additional evidence for the location of a core region of the 7A centromere, with a 5 Mb region of overlap between the two telosomes resulting from asymmetrical positioning of the breakpoints (Fig. 5A, Fig. 6). At the end of the 7AL telosome, evidence from the Bionano map indicated that the terminal 50Kb had been duplicated (in reverse complement) on the 7AL telosome, with this extended sequence not appearing in the 7AS side of the assembly. Coverage of raw 7AL read data across the IWGSC RefSeq v1.0 chromosome 7A centromere supports the presence of this sequence duplication at this end of the 7AL centromere (Fig. 6D, increased read coverage at centromere end of 7AL indicated by a dotted blue box); the duplication is absent from a standard chromosome 7A.

**Fig. 6.**
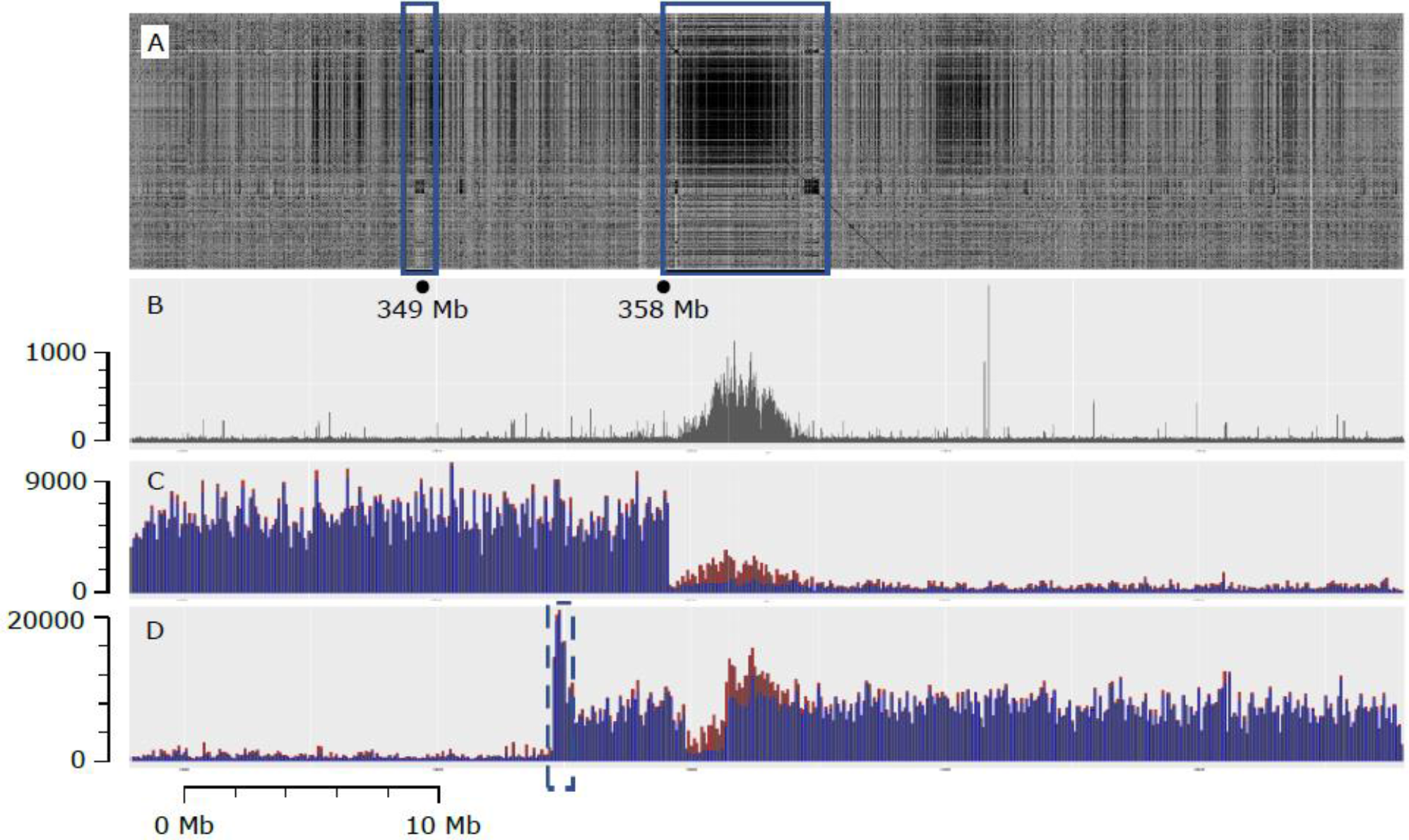
IWGSC RefSeq v1.0 chromosome 7A 338 Mb-388 Mb region. A. Dotplot of 338 Mb-388 Mb region against the 10Mb between 358 Mb and 368 Mb and indicates two regions (blue boxes) that are speculated to be integral to the centromere structure and involved in *in situ* CENH3 protein-antibody binding (Fig. S6); the left box at ca 349 Mb is suggested to have an incomplete genome assembly due to a breakdown in the assembly process as indicated in Fig. 5A (lower panel) since both the GYDLE and Bionano maps have breaks in the 349 Mb region. B. ChIP-Seq CENH3 data (SRA accessions SRR1686799 and SRR1686800) aligned to the 338 Mb-388 Mb region, counted in 10Kb bins. C. Raw CSS reads of 7AS (SRA accession SRR697723) aligned to the 338 Mb-388 Mb region (see also Additional file 8, Figure S7) 33 D. Raw CSS reads of 7AL (SRA accession SRR697675) aligned to the 338 Mb-388 Mb region (see also Additional file 8, Figure S7). The dotted blue box indicates a segment of the 7AL centromere that is duplicated as discussed in the text Unique alignments are shown in blue in both C and D and show the clear boundaries of 7AS and 7AL telosomes as well as a deletion in the 7AL telosome. Reads with multiple mapped locations are shown in red (single location selected randomly) and indicate that the core CRW region is represented in the raw 7AS reads, although at lower levels than on 7AL. Counts in bins of 100Kb.

The active centromere and associated kinetochore complex in plants can also be defined based on the location of the CENH3 binding domain (26). We aligned CENH3 ChIP-Seq data for wheat (25, 26) to the IWGSC RefSeq v1.0 and found a 5 Mb region on the proximal side of 7AL to the breakpoints (not in the region of overlap between the 7AS and 7AL assemblies) as the primary source of similarity to the CENH3 binding sequences and CRW repeat sequence families (Fig. 6A). This located the main CENH3 binding domain represented in the assembly to be on 7AL. Analysis of an independent assembly of Chinese variety Aikang 58 showed the same chromosomal structure, although the size of the core CENH3 binding/CRW repeat region was larger. Tiling of the GYDLE assembly around this region in IWGSC RefSeq v1.0 identified a gap in coverage of BAC data on the 7AL side of the assembly (Fig. 5A). Alignment of raw IWGSC CSS data across the region (Fig. 6C, 6D) showed a sharp drop in coverage to background levels at exactly the same location; however, alignment of the raw data used in the IWGSC RefSeq v1.0 assembly indicated this region was present in the whole-genome data (Additional file 8, Figure S7), implying a deletion of around 2Mb in the 7AL ditelosomic stock (Fig. 6).

Although the 7AS telosome appears to be missing a major CENH3 binding domain, records for tracking the transmission of the two telosomes in the Kansas Genetic Stock centre showed the transmission rates for the two telosomes were similar, implying that 7AS must also have an active centromere. We used the *in situ* localisation of the CENH3 antibody (Additional file 8 Figure S7, Additional file 13, (27)) to show that the 7AS telosome also has a localised CENH3 binding domain near the telosome breakpoint. Both telosomes carried a similar level of CENH3 antibody binding protein in the centromere regions, based on the analysis chromosome spreads shown in Additional file 8 (Figure S7), suggesting it unlikely that there exists a major difference in genome structure of the centromere. Furthermore, we found evidence that sequences from the Centromere Repeat of Wheat (CRW) region of this scaffold were present in the 7AS telosome, at low levels (Fig. 6A). Close inspection of the tiling of the GYDLE sequence around scaffold96327 (a single scaffold unconnected to the surrounding scaffolds in the pseudomolecule, also a single island in the GYDLE assembly) found highly dense copies of Byron CACTA elements (as well as representative copies of other CRW elements, Fig. 6A at position 349 Mb) and suggests this as a possible location for the 7AS CENH3 binding domain sequence within the 7A functional centromere region analogous to that found in rice centromere 8.

## Discussion

In this manuscript the resources for finishing a wheat reference genome sequence were defined at two levels, namely, micro-scale and macro-scale. At the macro-scale the IWGSC RefSeq v1.0 assembly provided a pseudomolecule against which our independent BAC-based assembly could be aligned, enabling a reduction in super-scaffold number, the completion of super-scaffold ordering and orientations and the local solving of micro-scale inconsistencies and deletions. This capacity enabled, across the whole chromosome, 52 CDS sequences in the IWGSC RefSeq v1.0 to have sections of Ns filled and gene models updated. In target regions, a method that combines multiple resources such as the raw CSS survey sequence (2), high density molecular genetic maps (28, Additional file 2) and Bionano maps was able to produce finished sequence (Methods, Additional file 3). The Bionano maps were particularly valuable as an independent source of linear sequence information when assemblies conflicted. Two target regions of chromosome 7A were studied in detail to explore the requirements for finishing the genome sequence of the reference assembly at a broader level. These sequences are the largest complete sequences available in wheat and highlight that merging sequences from multiple assemblies to achieve complete finishing is possible, but will require the re-referencing, preferably simultaneously rather than sequentially, of the multiple raw data sets and types to provide final validation where assembles agree, and to provide information to resolve conflicts between assemblies as these are found.

One of the 2.5 Mb regions that was finished overlapped the QTL initially defined by Huynh et al (16) for fructan content in the grain and in our analysis was shown to contain a tandem array of seven glycoside hydrolases (EC 3.2.1, labelled a to g) that were of particular interest since the gene model GH32b could be assigned to 1-FFT on sequence similarity basis and GH32g to 6-SFT. Both these genes are key in the fructan biosynthetic pathway (29). The GH32 genes were expressed in the grain and stem and the analysis of variation in grain fructan-levels from 900 wheat lines characterized using exome capture indicated that over half of the SNP variation in the QTL region associated with variation in grain fructan-levels located to the GH32 family genes. For the homoeologous GH32 array on chromosome 7D, the most highly significant association across the entire genome was also in this region and it is thus evident that selection at multiple loci is required for a phenotype such as grain fructan levels.

The 2.3 Mb region associated with TKW and spikelet number, within the broad yield QTL region on 7AL, required the more extensive integration of the IWGSC RefSeq v1.0 and GYDLE assemblies. Solving the complete sequence for this region showed distinct linkage blocks existed in diverse worldwide wheat lines, indicating that fine mapping this region through association analysis will be challenging. The gene families within linkage blocks included repetitive gene models annotated as housing domains involved in apoptosis as well as root morphology and thus provide targets for establishing a framework for strategies to select for variation which includes variation in copy number as suggested in (1).

The centromeres of chromosomes have been studied extensively (30) in microorganisms, animals and plants. The centromere of 7A was located within the C region (1) in chromosome 7A extending from position 240 to 410 Mb (170 Mb, (1)) and could be further defined as a 58 Mb region based on the presence of the reverse transcriptase sequence from the Cereba element (AY040832). Except for one unit located at 67 Mb in the telomeric region of 7AS, the Cereba element was unique to the centromere region within 7A. The detailed structure studies indicated that at least two domains for centromere activity existed within the functional domain that was syntenic to the rice chromosome 8 centromere. The centromere region contained 62 genes and 5 of these genes were also located in the rice chromosome 8 centromere and provided the basis for defining a syntenic functional centromere. Although the CENH3 binding sequences on 7AS were not as clearly defined as in 7AL, we speculate that this is due to a breakdown of the assembly process in the respective region (349 Mb region, see Fig. 5A, Fig 6A). The available data suggests the reduced CENH3 protein-antibody binding assayed in both the 7AS and 7AL telosomes (relative to the level of binding to normal chromosomes Additional file 8, Figure S6) is sufficient for the retention of centromeric activity. The analysis also indicated that the terminus of the 7AL centromere had a terminal 50Kb duplication of a sequence that is located between the two proposed CENH3 protein-antibody binding domains. In addition, an element, Tail (AB016967) (31) was found to have 100 units in the region 374.7 Mb – 376.9 Mb (on 7AL) and is unrelated to Cereba or the rice/maize centromere repeats but exists within the Quinta retrotransposable element. In situ hybridization (31) shows Tail is centromeric to all wheat chromosomes. The incursion of this most recent transposable element (Quinta/Tail) is a striking feature here in that the Tail sequence is a dispersed repeat in grasses related to wheat and is consistent with it being a recent addition to the wheat genome that has not had enough time to spread more widely. It is possible that new clusters of repetitive elements significantly enhance the network of interactions in which the centromere is involved in meiosis and mitosis (32).

## Conclusions

Chromosome 7A provided a useful model to carry out analyses that establish a foundation for developing an advanced, version 2.0, high-quality wheat reference genome assembly. The strategy developed in the present manuscript indicates that the required assembly algorithms and sequence data exist, while future investment in long read data, such as BioNano optical maps, will provide the complete resources necessary for integration of raw data into well-developed templates of the wheat reference genome, sufficient for the accurate interpretation of sequences from new wheat varieties. The suites of genes identified in regions of the genome associated with grain yield and quality provide a basis for identifying gene family copy number variation and new molecular markers for the rapid selection of difficult phenotypes in breeding programs. A key utility of the IWGSC RefSeq v1.0 genome assembly (1) is to accelerate QTL mapping and then support the gene cloning or perfect marker identification process in both fundamental and translational research. At the back end of these processes, it is the genome assembly quality that most often inhibits progress. Likewise, the use of gene editing and other similar modern breeding methods requires base level accuracy in focus regions. Importantly, the finished regions described in this research span the flanking markers of known QTL and hence these regions can be studied in full without unknown assembly issues impeding progress.

## Methods

### Independent assembly of chromosome 7A

The BAC library of 119,424 BACs (58,368 and 61,056 on 7AS and 7AL respectively) from flow-sorted chromosome arm 7A DNA was fingerprinted using the SNapShot method (19) and assembled into physical contigs using LTC software (20). The physical map comprised 732 BAC contigs and an MTP of 11,451 BACs totalling an estimated 755 Mb. For each physical contig, the MTP BACs were pooled into groups of no more than 20 BACs. These BAC pools were then shotgun sequenced using Illumina paired-end technology. The BAC pool sequence data were first assembled separately for each physical contig using Abyss, totalling 882 Mb in 74,572 contigs. The BAC pool based contigs, provided the starting point for integrating the various data sets using GYDLE software (Philippe Rigault, GYDLE Inc., Canada, https://www.GYDLE.com/bioinformatics; (38) (41)). An initial multiple alignment was produced using the NUCLEAR software (GYDLE Inc) as part of the hybrid assembly of the available datasets. Reprocessing of BAC pools assemblies identified BAC ends and removed low quality reads, and thus allowed BAC clones to be identified that were not true components of the respective pools. VISION software (GYDLE Inc) was used to visualize assemblies in a semi-manual curation process with assembly metrics calculated using Perl, R and Shell scripts. An iterative process provided the basis for integrating extensive mate-pair data, Bionano data and Bayer-IWGSC Whole Genome Profiling (WGP^TM^) tags*(1)*. The three stages can be summarized as; 1) integrating the BAC pool mapping and sequencing data with multiple mate-pair datasets (see also Additional file 1); 2) extending and refining scaffolds based on iterative realignments of the sequence data; 3) cross-validating the sequence assembly with physical mapping data to link scaffolds with physical contigs, identifying missing BACs, contaminations and physical contig errors, and allowing for selected regions to undergo interactive editing and visualisation in order to produce locally finished, manually-reviewed sequence. It was possible to connect consecutive BAC pool sequence assemblies using Bionano optical maps generated from flow sorted Chinese Spring 7AS/7AL telosomic lines with the sequence structure visualized by fluorescent labelling of Nt.BspQI nickase (GCTCTTC) sites (details below), to construct 124 scaffolds or “islands” covering 735.1 Mb. The 18 largest islands comprised more than 50% of the total sequence.

The GYDLE website (https://www.GYDLE.com/) provides information on accessing the software as well as the solutions and services provided by the GYDLE company. The scale and novelty of this work required not only capabilities that were (and still are) not available in any other product (open source or commercial), but also specific developments to accommodate both the integration of specific data and their visualisation (e.g. Figures 1, 2B, 3A, 3B). The GYDLE software NUCLEAR and VISION have been utilized in the analysis of several genomes, including the Eucalypt (38) and wheat genomes (1)(41).

### BAC library fingerprinting

The BAC clones 7A BAC MTP were fingerprinted as described in (19). The use of an ABI3730XL, with a more sensitive laser improved fingerprinting resolution and made it possible to reduce the amount of BAC DNA sample for electrophoresis, thus lowering fingerprinting costs. 0.5-1.2 μg instead of 1.0-2.0 μg of BAC DNA were simultaneously digested with 2.0 units each BamHI, EcoRI, XbaI, XhoI, and HaeIII (New England Biolabs, Beverly, Massachusetts) at 37C for 3 hrs. DNAs were labeled using the SNaPshot kit (0.4 μl of reagent, Applied Biosystems, Foster City, California) at 65C for 1 hr and precipitated with ethanol. DNAs were dissolved in 9.9 μl of Hi-Di formamide, and 0.3 μl of Liz1200 size standard was added to each sample. Restriction fragments were sized on the ABI3730XL. Raw outputs from BAC fingerprinting were converted to .gm format using GeneMapper and filtered with Genoprofiler. Resulting files consisted of lists of numbers denoting fragment size for each BAC, added to an offset for each colour: 0 for blue, 10,000 for green, 20,000 for yellow, 30,000 for red.

### Sequencing of MTP BACs

BAC clone DNA was prepared by standard alkaline lysis mini-prep procedure. BAC clones were grown overnight 1.2 ml of 2YT media with chloramphenicol in 96-well culture plates. Plates were spun by centrifugation at 2500g for 10 mins to pellet cells. Each pellet was resuspended in 400 µl of GTE buffer (0.05M glucose, 0.01M EDTA, 0.025M Tris pH 7.4). 60µL of the resuspended cells were transferred to an extraction plate, and 100µL of NaOH/SDS solution (0.8%NaOH, 1%SDS) was added to lyse the cells. This solution was neutralized by the addition of 100µL of potassium acetate (3M), and gently mixed by inversion. Lysates were vacuum-filtered through a Costar 96-well Filter plate (0.2µm GHP) and precipitated by the addition of 110µl isopropanol. BAC DNA was pelleted by centrifugation at 2500g for 15 minutes. Supernatant was removed, and the pellets washed once with 200µl of ice-cold 70% ethanol. The pellet was let air-dry for 20-30mins and resuspended in 50µL of water.

### Illumina sample Preparation and Sequencing

100 ng of BAC DNA was sheared in 50 µl by ultra-sonication (Covaris E220 instrument settings Duty Factor = 5%, Intensity= 5, Cycles per burst = 200, Duration = 55 sec, Displayed Power 13 W, temperature 5.5-6.0 °C). Samples were processed using the Illumina TruSeq HT DNA sample preparation kit (FC-121-2003) as per the manufacturer’s guidelines. Following ligation of adapters, a “double-sided” SPRI size selection was performed to select for library fragment with a median size of 550-600 bp. Libraries were assessed by gel electrophoresis (Agilent D1000 ScreenTape Assay, Cat. No. 5067-5582 and 5067-5583) and quantified by qPCR (KAPA Library Quantification Kits for Illumina, Cat. No. KK4835). Sequencing was performed on the HiSeq 2500 system using TruSeq Rapid PE Cluster Kit HS (Cat. No. PE-402-4001) and TruSeq Rapid SBS Kit HS (Cat. No. FC-402-4001 and FC-402-4002).

The minimum tiling paths (MTPs) of contigs from the first version of the physical assembly were used to define pools of BACs for sequencing. Large pools (over 20 BACs in MTP) were split into multiple pools. 100 ng of pooled BAC DNA was fragmented by ultra-sonication (Covaris E200, Covaris Woburn, MA USA) and DNA libraries with an insert size of 450 bp were prepared using Illumina TruSeq DNA HT Sample Preparation Kit (Illumina San Diego, CA, USA). The size of each library was validated using the DNA 1000 ScreenTape (Agilent Santa Clara, CA, USA) and quantified by qPCR before normalization and pooling. 96 BAC pool libraries were sequenced in one lane of the Illumina HiSeq 2500 in rapid mode with 2×150 bp paired end reads (Illumina San Diego, CA, USA).

### Read filtering and removal of bacterial sequences

All available *E. coli* genome sequences in NCBI were used to remove non-wheat sequences because some sequences were found from unexpected strains. The reads were QC’d to remove contaminating sequences and poor-quality reads before running assembly scripts.

### Mate-pair sequencing

Amplified DNA was produced from the DNA isolated from flow-sorted 7AS and 7AL telosomic chromosome arms using flow-sorted chromosomes treated with proteinase K and amplified using Phi29 multiple displacement amplification (MDA). Overnight amplification in a 20-microlitre reaction produced 3.7 – 5.7 micrograms DNA with a majority of products between 5 and 30 Kb. This amplified DNA was then processed to remove nicks and single-stranded DNA before carrying out the Nextera Mate Pair / HiSeq System (following manufacturer’s instructions) for generating a high coverage of mate-pair sequence information. The libraries covered 200 - 5000 bp.

### PacBio sequencing

Short-read data and PacBio sequencing of a single BAC (7AS-066B03) followed protocols provided by the technology provider.

### Bionano view of genome sequence

A total of 2.8 million of each of the 7A arms, corresponding to 1.14 ug DNA, were purified by flow cytometric sorting as described above with purities of 80%, and 86% for the 7AS and 7AL arm, respectively. Chromosome arm DNA was used to construct Bionano maps following the protocol of Staňková et al. (17). Based on the frequency of recognition sites in the survey sequences of 7A arms (IWGSC, 2014), Nt.BspQI nickase (GCTCTTC recognition site) with estimated frequency of 11 sites/100 Kb was selected for DNA labeling. Chromosome arm DNA samples were labeled at nicking sites with Alexa546-dUTP fluorochrome, their DNA was stained with YOYO and analyzed on the IRYS platform (Bionano Genomics, San Diego, USA). Bionano maps of 7AS and 7AL, assembled *de novo* using molecules longer than 150 Kb, exhibited coverage of 192x (79 Gbp) and 238x (97 Gbp), respectively.

*De novo* assembly of Bionano maps was performed by a pairwise comparison of all single molecules and graph building (37). A p-value threshold of 1e-^10^ was used during the pairwise assembly, 10e^−11^ for extension and refinement steps, and 1e^−15^ for a final refinement. The use of Bionano data in the 7A assembly is a significant advance over the work of Stankova et al (2016), as the GYDLE software performs a scalable and true hybrid optical/sequence assembly enabling local sequence resolution (e.g. gaps, tandem repeats) based on systematic comparisons of distances in optical and sequence space, as well as map validation using molecule data.

### Linkage disequilibrium analysis

A diverse spring bread wheat collection (n=863) comprising of landraces and elite cultivars was used in this study to understand the haplotype structure and extent of linkage disequilibrium (LD) in the yield QTL region on 7A, coordinates 671200000-675300000 bp. LD values were estimated and visualized using the Haploview software (42) and only common SNPs with high minor allele frequency (MAF > 0.3) and present within 2000 bp on either side of gene were included in this analysis. A total of 203 SNPs within 35 gene models (plus 2000 bp on either side) spanning the whole region were identified. We could not detect any common SNPs in the remaining 18 genes in the QTL region. The associations (Fig. 4) were color coded as: bright red D’ =1.0 and LOD ≥ 2.0 (high LD); light shades of red indicate D’ < 1.0 and LOD ≥ 2.0 (low-medium LD); white indicates D’ <1.0 and LOD <2.0 (no LD or complete decay).

### Defining the centromere

To confirm the presence of a large missing CENH3 binding domain in the 7AS di-telosomic stock we aligned the 7AS (SRR697699, SRR697706, SRR697723) and 7AL (SRR697675, SRR697676, SRR697680), 101 bp paired-end Illumina reads generated for the CSS assembly to the chromosome 7A wheat-AK58 assembly using NUCLEAR (GYDLE) with filtering for minimum base quality of phred 20, minimum length per side of 50 and paired reads only, and mapping parameters allowing a single mismatch in an HSP of length 50, a minimum alignment length of 50 bp, a sensitivity of 25 and a k of 13 (~98% identity). See also Additional files 12 and 13.

## Availability of data and material

- Wheat chromosome 7A mate-pair data from flow-sorted chromosomes (45).
- IWGSC Wheat Chromosome 7A BACs sequenced in pools based on the physical map Minimum Tiling Path (MTP) with Illumina HiSeq 2500 (46).
- Sequencing of a Chinese spring wheat with 7EL addition from Thinopyrum elongatum (47).
- Finished regions of chromosome 7A in fasta format and Bionano assemblies (48).

## Abbreviations

Contig: consensus region of DNA sequence represented by overlapping sequence reads. Can have unresolved bases (N), but no gaps.
Scaffold: consensus region of DNA sequence represented by ordered (but not necessarily oriented) contigs, separated by gaps of known (estimated) length.
Island: genomic region represented by overlapping sets of DNA sequences (scaffolds), physical entities (optical map or molecule, physical clone), or both.
Super-scaffold: a portion of the genome sequence where scaffolds have been ordered and oriented relative to each other

## Declarations

- <u>Ethics approval and consent to participate</u>: Not required.
- <u>Consent for publication</u>: Authors have provided consent.
- <u>Availability of data and material</u>: The new genome sequence data is submitted to the IWGSC Data Repository hosted at URGI: https://wheat-urgi.versailles.inra.fr/Seq-Repository
- <u>Competing interests</u>: PR, SB, and M-AN have competing commercial interests as employees and stockholders of GYDLE, which is a commercial company that provides bioinformatics analysis software and services. This does not alter the authors’ adherence to all of the Genome Biology policies on sharing data and materials. The remaining authors declare that they have no competing interests.
- <u>Funding</u>: Australian Government Department of Industry, Innovation, Science, Research and Tertiary Education (funding agreement ACSRF00542), BioPlatforms Australia (BPA) and Grain Research Development Corporation (agreement UMU00037) are thanked for funding the chromosome 7A project. CSIRO Plant Industry, Australia, funded the establishment of the MAGIC molecular genetic map. Agriculture Victoria Research funded bioinformatics capacity and infrastructure. The CENH3 antibody/cytological studies were supported by NSF grant contract 1338897. The work of FC was supported by the INB ("Instituto National de Bioinformatica") Project PT13/0001/0021 (ISCIII-FEDER). Chromosome flow-sorting, construction of BAC libraries and Bionano maps were partially supported by Czech Ministry of Education Youth and Sports (award LO1204 from the National Program of Sustainability).
- <u>Acknowledgements</u>: The authors are grateful to DM Appels for her dedication in establishing the early molecular genetic maps for chromosome 7A. Bernd Friebe is acknowledged for his guidance in the cytological studies of 7AS and 7AL. Part of this work was supported by resources provided by the Pawsey Supercomputing Centre with funding from the Australian Government and the Government of Western Australia. The authors are grateful to Prof Jia Jizeng for discussion in relation to the genome assembly of Aikang58.
- <u>Authors’ contributions</u>: GK-G, PR, JT contributed equally to experimental design, data analysis and interpretation/writing of manuscript;, RP, MH, KF, RA genome analyses and interpretation; ZF, AK data analysis and physical map construction; EH, CC, JT MAGIC map construction; MA rice-wheat phylogenomic; AS, DK, mate-pair libraries; PS. BD, FC, PL, Chinese Spring x Renan molecular genetic map and annotation of genome sequence; NW-H, UB, PE, DF, AJ analysis of gene space and QTL, SB, M-AN, development of Bionano alignments and tools; JD, HŠ, JŠ, HT, flow sorting of chromosomes, BAC library construction and Bionano maps; M-CL fingerprinting of BAC library; FC, MP gene annotation; DC, ZR, AK58 genome assembly, JN-P, genome assembly; DI in-house software development; IWGSC, genome assembly network, D-HK, CENH3 antibody cytology; RA, planning of experiments and writing of manuscript.

## Additional files

**Additional file 1:** BAC preparation and analysis for physical maps, Microsoft Word document .docx, 17KB

**Additional file 2a:** Combining the MAGIC 8-way cross 7A and Chinese Spring x Renan 7A maps, Microsoft Word document .docx, 18KB

**Additional file 2b:** Curated genetic map of 7A for anchoring the genome sequence, Microsoft Excel Worksheet .xlsx, 115KB

**Additional file 3:** DNA sequence assembly details, Microsoft Word document .docx, 16KB

**Additional file 4a:** IWGSC RefSeq 7A scaffolds related to PacBio and GYDLE scaffolds, Microsoft Excel Worksheet .xlsx, 18KB

**Additional file 4b:** GYDLE island IDs related to Bionano maps and BAC sets, Microsoft Excel Worksheet .xlsx, 38KB

**Additional file 5:** IWGSC RefSeq v1.0 7A related to GYDLE scaffolds and contigs plus bridging sequences, Microsoft Excel Worksheet .xlsx, 398KB

**Additional file 6:** IWGSC RefSeq v1.0 7A gene IDs corrected by manual curation, Microsoft Excel Worksheet .xlsx, 21KB

**Additional file 7:** Genome association analyses for variation in grain fructan and yield (grain number), Microsoft Word document .docx, 17KB

**Additional file 8:** Figures S1 – S7 (with legends) for Additional files, Microsoft Word document .docx, 5,814KB

**Additional file 9:** 7A QTL analysis and coordinates in RAC875 x Kukri cross, Microsoft Excel Worksheet .xlsx, 24KB

**Additional file 10:** Finished 7A yield region assembly BAC sets, Microsoft Excel Worksheet .xlsx, 11KB

**Additional file 11:** 7A yield region annotation, Microsoft Excel Worksheet .xlsx, 21KB

**Additional file 12:** Rice chromosome 8 centromere synteny analysis with chromosome 7A, Microsoft Word document .docx, 15KB

**Additional file 13:** Quantification of centromere fluorescence for CEN H3 antibody in situ locations, Microsoft Word document .docx, 15KB

